# The Fruit Fly Brain Observatory: From Structure to Function

**DOI:** 10.1101/580290

**Authors:** Nikul H. Ukani, Chung-Heng Yeh, Adam Tomkins, Yiyin Zhou, Dorian Florescu, Carlos Luna Ortiz, Yu-Chi Huang, Cheng-Te Wang, Mehmet K. Turkcan, Tingkai Liu, Paul Richmond, Chung-Chuan Lo, Daniel Coca, Ann-Shyn Chiang, Aurel A. Lazar

## Abstract

The fruit fly is a key model organism for studying the activity of interconnected brain circuits. A large scattered global research community of neurobiologists and neurogeneticists, computational and theoretical neuroscientists, and computer scientists and engineers has been developing a vast trove of experimental and modeling data that has yet to be distilled into new knowledge and understanding of the functional logic of the brain. Developing open shared models, modelling tools and data repositories that can be accessed from anywhere in the world is the necessary engine for accelerating our understanding of how the brain works.

To that end we developed the Fruit Fly Brain Observatory (FFBO), the next generation open-source platform to support open, collaborative Drosophila neuroscience research. FFBO provides a (i) hub for storing and integrating fruit fly brain research data from multiple data sources worldwide, (ii) unified repository of tools and methods to build, emulate and compare fruit fly brain models in health and disease, and (iii) an open framework for fruit fly brain data processing and model execution. FFBO provides access to application tools for visualizing, configuring, simulating and analyzing computational models of brain circuits of the (i) cell type map, (ii) connectome, (iii) synaptome, and (iv) activity map using intuitive queries in plain English. Tools are provided to extract the function inherent in these structural maps. All applications can be accessed with any modern browser.

## Introduction

Animal behavior is governed by the activity of interconnected brain circuits. Comprehensive brain wiring maps are needed to formulate hypotheses about information flow and also to guide genetic manipulations aimed at understanding how genes and circuits orchestrate complex behaviors. The availability of a powerful toolbox of transgenic methods for neuronal circuit analysis, wide variety of mutants, the ease of culture and short development cycles makes *Drosophila* an ideal model system for investigating the relationship between genes, brain structure, function and behavior [1–3].

A successful determination of how the brain’s highly complex structure implements specific functions requires its decomposition into functional modules whose input-output relationships can be individually analyzed and whose interactions can be explained in terms of the groups of synaptic connections that exist between them. Understanding how the fruit fly brain works, requires the assembling of four major reference maps, namely, (i) the cell type map - classification of cells into groups characterized by their morphology, molecular expression pattern and distribution/innervation in all brain regions; (ii) the connectome - the wiring diagram of the nerve connections among all neurons in the brain; (iii) the synaptome - the set of all synapses, their distribution and expressed neurotransmitter and receptor type in all brain regions, and (iv) the activity map - the electrical activity of all neurons in the brain associated with a particular brain state.

In the last century pioneering work began to map out these four areas on a small scale. For example, staining-based cell type classification has led to the FlyBrain, the very first online atlas of the *Drosophila* brain [4]. Around the same time, Flybase was created as a database of *Drosophila* genes and genomes [5]. Electron Microscopy (EM) has been used to precisely map columnar elements in the lamina neuropil [6]. Electrophysiological recordings of neurons such as Lobula Plate Tangential Cells has played a central role in gaining insight into visual motion detection [7].

In the past decade, biological data contributing to the cell type map, connectome and synaptome of the fruit fly brain have been rapidly and increasingly made available thanks to advancing genetic technology and volume EM technology. A number of genetics data libraries have been greatly expanded, including the FlyBase [8], and others newly created. For example, functional genomics data is provided by the Drosophila RNAi Screening Center (DRSC) and Transgenic RNAi Project (TRiP) [9], RNAi Libraries by the Vienna Drosophila Research Center [10] and FlyLight GAL4 and Split-GAL4 lines by the HHMI Janelia Research Campus [11, 12]. Furthermore, EM-based data has been provided for the lamina cartridge [13], medulla [14, 15] and mushroom body [16]. Most recently, the whole fly brain has been imaged using EM [17]. In addition, the cell type map of the mushroom body [12], medulla [15] and central complex [18, 19] have been published. A large mesoscale connectome of the fruit fly brain has been provided (FlyCircuit) [20] and the connectome of 7 medulla columns (Janelia) [15] has been reconstructed, and made publicly available. In contrast, despite the strong interest in *Drosophila* neuron recordings, open neurophysiological data can rarely be found in the public domain. One of the exceptions is the DoOR database for mapping *Drosophila* odorant responses [21].

This diverse set of data has been utilized by three stakeholder groups towards understanding the function of fly brain circuits (see Figure 1). The first group consists of neurobiologists and neurogenetists who are interested to find neurons, genes, genetic lines to perform experiments. Many existing fruit fly brain databases, web services and software, such as FlyCircuit DB [20], FlyBrain Neuron Database [22], Virtual Fly Brain [23] and BrainBase [24], serve this group of researchers. They enable search and visualization of confocal imaging stacks of lines/neurons and/or information describing individual neurons.

**Figure 1:**
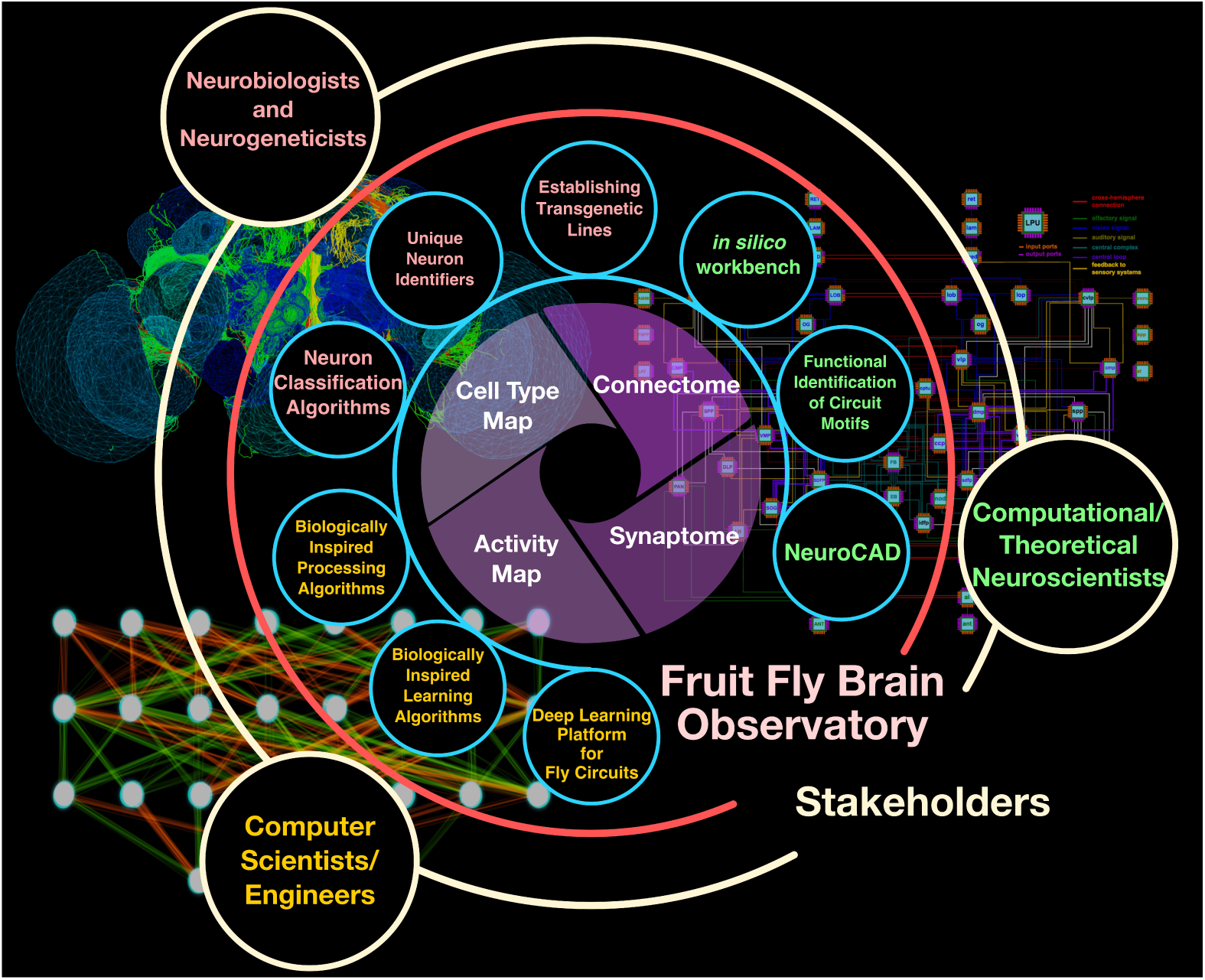
The FFBO open collaborative ecosystem. The FFBO assembles the cell type, connectome, synaptome and activity maps within a single ecosystem that benefit three group of stakeholders, namely, the neurobiologists/neurogenetists, the computational/theoretical neuroscientists and the computer scientists/engineers. The FFBO will, in the future, build additional tools, such as those in the small circles shown in the middle ring, to help each group of stakeholders to contribute to building the 4 maps as well as to fully utilize them for understanding the function of the fly brain.

The second stakeholder group consists of computational/theoretical neuroscientists. Several computational models have been proposed to study whole brain function [25], the early visual system [26] and visual motion detection [27, 28]. Electrophysiology data has been used to identify the computation underlying photoreceptor function [29]. A number of recent computational investigations addressed the function of the central complex (CX) [30–33] whose anatomical structure, physiological properties and behavioral connection have been subject to active recent studies [18, 19, 34, 35].

A third stakeholder group consists of computer scientists/engineers who independently investigated pattern classification using deep learning and/or applied such methods to model the lamina cartridges and medulla columns of the fly [36, 37]. The overriding goal here is to create novel circuits and models for deep learning [38] and reinforcement learning [39].

While extensive amounts of data has become increasingly influential within the domain of each of the stakeholders, researchers are still faced with several challenges including the lack of a (i) hub for storing and integrating fruit fly brain research data from multiple data sources worldwide, (ii) standardized repository of tools and methods to build, emulate and compare fruit fly brain models in health and disease, and (iii) open framework for fruit fly brain data processing and model execution.

The Fruit Fly Brain Observatory (FFBO) presented here addresses these challenges heads-on (see Figure 1). FFBO provides all stakeholders with the (i) means to build and access the four maps (see inner circle), and (ii) tools to build upon these maps the algorithms tailored to their respective knowledge domain (some examples are briefly described in the circles in blue surrounding the inner circle).

## Results

### The Fruit Fly Brain Observatory

The FFBO assembles the cell-type, connectome, synaptome and activity maps within a single ecosystem, and more importantly, supports the building of the functional map of the fruit fly brain. The latter map is a key step in gaining insights into the functional logic of the fruit fly brain circuits and their I/O behavior at different levels of abstraction.

The system architecture of the FFBO (shown in Figure S1) is built around an expanding modular architecture. At the heart of the FFBO is the NeuroArch database [40], that integrates both biological data and computational models of brain circuits into a single database, and Neurokernel, a GPU-enabled computational engine for emulating the fruit fly brain [41]. The integrated database can be leveraged through the NeuroNLP and NeuroGFX front-ends. NeuroNLP enables researchers to query biological data in plain English, including morphology and position of neurons (cell type map), connectivity between neurons (connectome) and distribution and type of synapses (synaptome). Moreover, it provides the first open neurophysiology data service for the fruit fly brain (activity map). NeuroGFX offers users means to explore the functional map of the fly brain circuits by providing them with a highly intuitive graphical interface to configure, compose and execute neural circuit models within Neurokernel. The main modules of the FFBO architecture are briefly described below.

**NeuroArch** is a graph database for codifying knowledge about fruit fly brain circuits. It is designed with two user communities in mind: (i) neurobiologists/neurogeneticists interested in querying the database to address questions regarding neuroanatomy, neural circuits, neurons, synapses, neurotransmitters, and gene expression, and (ii) computational/theoretical neuroscientists and computer science/engineers interested in the instantiation of models of neural circuits and architectures, their program execution, and validation of hypotheses regarding brain function. A key aim of NeuroArch is to provide a resource that supports and connects the research carried out by these two communities. To this end, NeuroArch defines a data model for representation of both biological data and model structure and the relationships between them within a single graph database [40].

The connectomic and anatomical data currently in the FFBO platform includes all available open fly brain data from the (i) FlyCircuit [20], spanning some 20,000 neurons and 1,260,000 inferred synaptic connections [25], and (ii) the Janelia seven column Medulla EM reconstruction that includes some 500 neurons and 67,000 synapses [15, 42], and (iii) the Janelia larval EM reconstruction that currently includes some 500 neurons and 138,000 synapses [43, 44]. The physiological data currently in the database consists of 1.6 hours of electrophysiology recordings from photoreceptors [29, 45], olfactory sensory neurons [46] and antennal lobe projection neurons [47].

The current NeuroArch database also includes two different models of the retina, developed by two research groups, and a model of the lamina neuropil of the Drosophila. Additionally, a model of the early olfactory system, including the antenna and antennal lobe resides in the current NeuroArch.

**Neurokernel** is an open-source engine implemented in Python for the collaborative emulation and validation of fruit fly brain models on multiple Graphics Processing Units (GPUs) [41]. Neurokernel provides a programming model based on the structural organization of the fly brain that consists of some 50 functional modules called Local Processing Units (LPUs) and the connectivity patterns that link them. Neurokernel defines application programming interfaces for communication between LPUs regardless of their internal design. Researchers can independently model different regions of the fly brain as LPUs and easily interconnect these for more complex functional validations.

**NeuroNLP** provides a modern web-based portal for navigating biological data relating to fruit fly brain circuits. It is equipped with a user-friendly, graphical interface to aggregate cell-type, connectome, synaptome and physiology data in the NeuroArch database, with the ability to simultaneously query against and retrieve information from disparate datasets.

NeuroNLP features a novel *natural language interface* that constructs complex queries against the underlying database from plain English instructions such as “*show GABAergic neurons that have dendrites in left antennal lobe and axons in both left lateral horn and right dorsolateral protocerebrum*” (or simply “*show GABAergic neurons that have dendrites in al and axons in both lh and DLP*”). This provides highly intuitive access to the integrated fruit fly brain circuit data, without the presumption of knowledge of a query language, syntax or cumbersome user interfaces. The results of the queries are presented using powerful 3D visualization and can be shared using a tag (see Methods, Figure S2 and Video S1) or by a demo script (see Methods and Video S2) for publication and collaboration. In addition, any neuron in the scene can be explored in greater detail using the information panel, which provides a one stop access to all data associated with a particular neuron (see Methods).

**NeuroGFX** provides an environment to easily explore circuit structure and function ulti-mately leading to biological validation. On the whole brain level NeuroGFX lays out the guidelines for the development of whole brain emulation. On the neuropil level, NeuroGFX allows users to study the I/O of each LPU. The canonical circuits (circuit motifs) are also identified on this level and NeuroGFX can be used to study the effect of different compositions mediated by local neurons.

NeuroGFX features a set of highly intuitive tools for exploring the function of neural circuit models, which can be accessed through a graphical user interface (GUI), allowing the user to (i) associate circuit diagrams with biological data, (ii) graphically construct an *in silico* experiment and execute manipulated circuits on GPUs, (iii) visualize the execution results in the context of biological brain structure. These capabilities are supported by a seamless integration of the NeuroArch database and the Neurokernel engine in the FFBO architecture.

Specifically, NeuroGFX assembles executable circuits of the fruit fly brain neuropils through composition of queries of the NeuroArch database that hosts executable models alongside biological data (for an example, see Methods). The resulting circuits are optimized for parallel execution by the Neurokernel Engine on general purpose Graphics Processing Units (GPGPUs) [41]. NeuroGFX enables the exploration of the function of the fruit fly brain on 3 levels of abstraction on (i) the whole brain level, (ii) the neuropil level and (iii) the local circuit level. On all levels, the goal of NeuroGFX is to help decipher the mechanisms underlying the function of circuits extracted from biological data through the construction, manipulation and comparison of executable brain circuits.

### Exploring Fruit Fly Brain MAPs with NeuroNLP

#### Building the Cell Type Map

NeuroNLP enables the study of cell types and the construction of circuits in the cell type map. As an example, with the simple query “*show Mi1 neuron in the home column in Medulla*”, the Mi1 neuron that belongs to the home column in the Janelia Medulla dataset can be visualized (see Figure 2a far left in orange, and also Video S3). The majority of the neuron morphology data do not have assigned cell types, however. NeuronNLP still allows the user to probe certain neurons known in the literature. For example, if the innervation pattern of such cells is known, the following queries can be used: “*show neurons that connect neuropil A to neuropil B*” or “*show neurons that have a dendrite in neuropil A and an axon in neuropil B*”. Queries can also be used to filter or modify the current set of results, for example using “*keep/remove cholinergic neurons*”, it is possible to filter neurons by the expression pattern of their neurotransmitter. [48] has characterized over 20 types of Lobula Plate Tangential Cells (LPTCs) in the blowfly. Using NeuroNLP, 8 types of LPTCs are revealed by a series of queries in NeuroNLP, as shown in Figure 2b. 6 out of them each corresponds to a blowfly LPTC type, but the innervation pattern of the other two have not been previously described (see Video S4).

**Figure 2:**
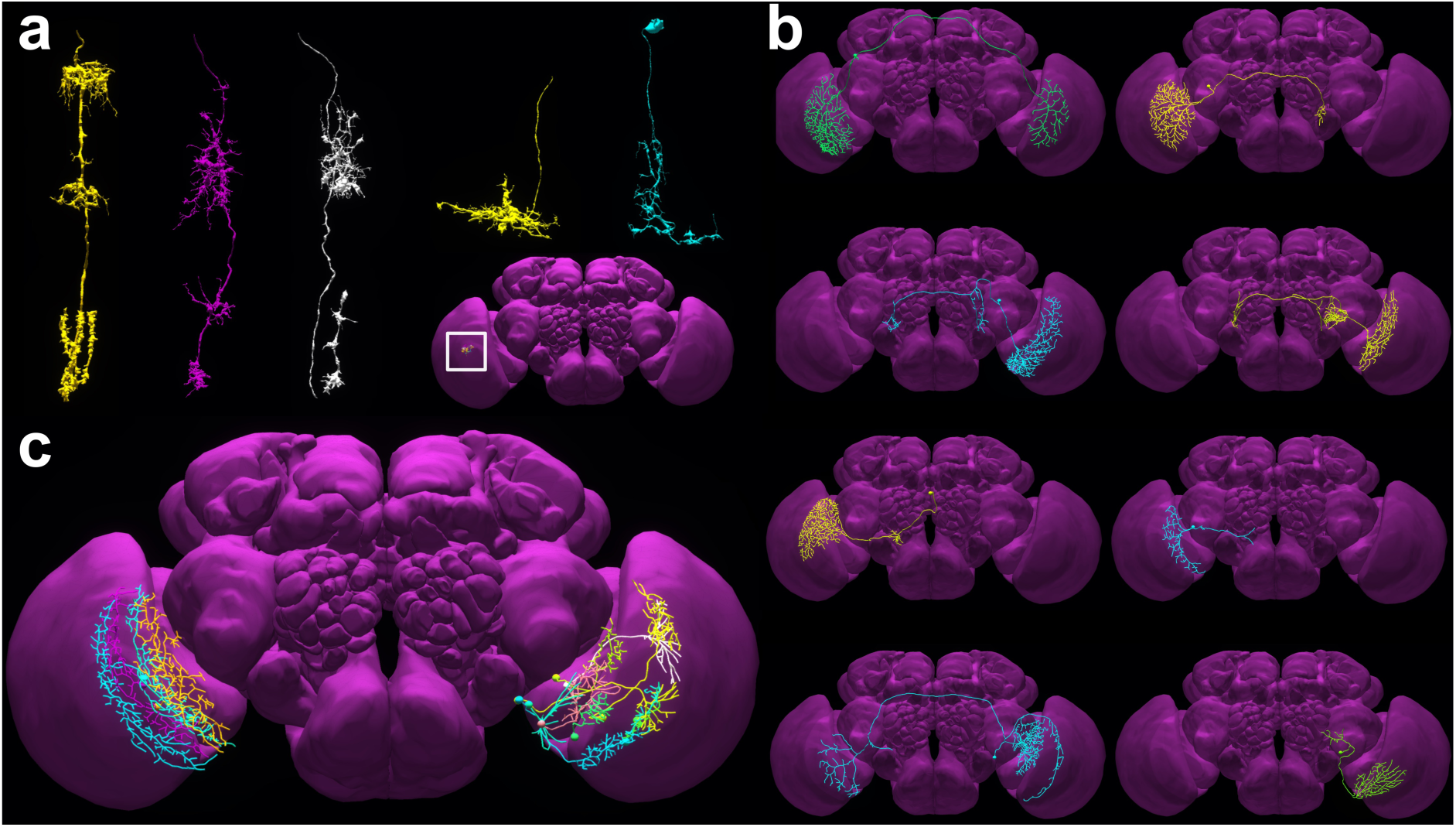
Querying from and building a cell-type map in FFBO. FFBO integrates different fly brain data sources to provide a queryable cell type map. The wealth of neurons in the database can be used to expand the current known cell-type map. **(a)** Neurons of existing cell types can be directly queried by their name, e.g., “Show Mi1 neuron in the home column”. Exemplary neurons are shown, from left to right, Mi1, Mi4, Mi9, Dm8 and Dm2. Inset indicates the approximate location of these neurons in the brain (see also Video S3). They can be accessed with the tag “paper:fig2a_v2”. **(b)** 8 types of lobula plate tangential cells (LPTCs) revealed by a series of queries in NeuroNLP. 6 of them each has a homology in calliphora that was previously described in [48], and two of the shown types have no obvious homology in the previous literature. With queries of presynaptic and postsynaptic partners, it is possible to obtain, respectively, tentative circuits that give rise to the function of the LPTC and subsequent circuits that further process information encoded in the LPTC (see also Video S4). Access in NeuroNLP for: H1 with tag “LPTC-H1”, H2 with tag “LPTC-H2”, H3 with tag “LPTC-H3”, H4 with tag “LPTC-H4”, CH with tag “LPTC-CH”, VS with tag “LPTC-VS”, Unknown type 1 with tag “LPTC-unknown1”, and Unknown type 2 with tag “LPTC-unknown2”. **(c)** Queries in NeuroNLP also reveal a subset of Lobula Plate intrinsic (LPi) neurons of the Lobulat Plate (on the left in the right Lobula Plate, see also in NeuroNLP with tag “glutamatergic_LPi_v1”), that has a different tiling pattern from those of the LPi neurons that innervates layers 3 and 4 as described in [49] (on the right in the left Lobula Plate). Such a tiling pattern may suggest that these neurons may innervate layer 1 and 2 of Lobula Plate and interact with the T4 neurons that encode horizontal motion in a similar way to the LPi neurons in layer 3/4 that interact with T5 neurons (see also Video S5).

In addition to finding previously known cell types, NeuroNLP supports the search for new cell types. In Figure 2c, the NeuroNLP query reveals a new type of Lobula Plate intrinsic (LPi) neurons that innervate layers 1 and 2 of the Lobulat Plate. These neurons are the sister neurons of the LPi neurons that innervate layers 3 and 4 in [49] (see Video S5).

#### Building the Connectome

NeuroNLP can be used to query the connectivity between fly brain neurons. The resulting information is critical for constructing and exploring model brain circuits and pathways. Brain circuits can be built up in the NeuroNLP workspace, by combining results from successive queries, for example “*add GABAergic neurons in EB*” or “*add postsynaptic neurons*”. Alternatively, individual presynaptic/postsynaptic partners can be added to the scene through the information panel. The connectivity of the resulting neural circuit can be exported from the GUI to a CSV file.

In Figure 3a-c, we demonstrate how to build a neural circuit responsible for visual motion detection. The query about connectivity is highly intuitive. We start from a T4a neuron (Figure 3a) that is known to be directional selective to front-to-back motion, and trace back to the neurons that provide direct inputs to it by adding presynaptic neurons with at least 3 synapses (Figure 3b), and those that provide indirect inputs by adding again presynaptic neurons with at least 5 synapses from the result of the first query (Figure 3c). This allows us to easily build a motion pathway and notice that not only columnar neurons but also a large number of non-columnar ones may play a critical role in motion detection (see Video S6).

**Figure 3:**
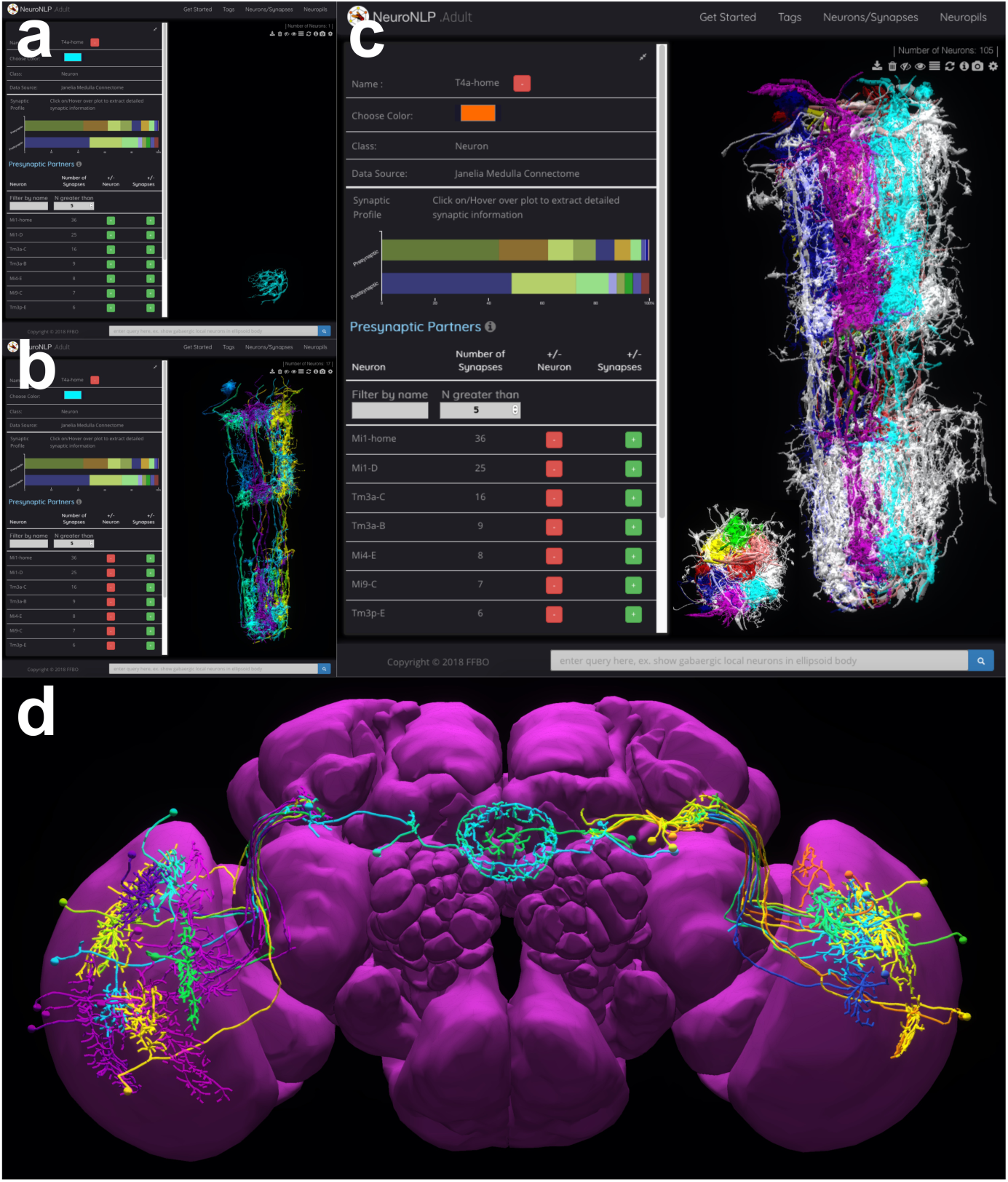
Constructing neural pathways based on the connectivity map in NeuroNLP. **(a-c)** Querying neurons in the visual motion detection pathway. Starting from a T4a neuron (a) with tag “ffbo:fig3a_v1”, that is known to be directionally selective to an ON motion signal, we query a pathway in medulla that provides direct and indirect inputs to this T4a neuron. (b) Adding presynaptic neuron to this T4a neuron with at least 3 synapses provides the neurons that directly synapse onto the T4a neuron (tag: “ffbo:fig3b_v1”). (c) Adding presynaptic neurons to the neurons in (b) with at least 5 synapses expands the set of neurons that provide indirect inputs to the T4a neuron. Columnar neurons in columns home and A-F are colored with red, green, yellow, blue, magenta, cyan and brown, respectively. In addition to the columnar neurons, a large number of non-columnar neurons (white) are involved in the ON motion pathway (see also Video S6). Inset: Top-down view. Access in NeuroNLP with tag: “ffbo:fig3c_v1”. **(d)** Construction of a visual pathway into EB as described in [50] using the connectivity map. Neurons from Medulla directly innervate the lower part of the OPTU and make synaptic contact with the TB neurons. The TB neurons project to the lateral triangle where ring neurons have their dendrites.(see also Video S7). Access in NeuroNLP with tag “10.1038/nn.4581_v0.1”.

In [50], a two-stage visual pathway between the medulla and the bulb (BU, or lateral triangle) was uncovered. This identifies potential sources of visual input to ring neurons that link the BU and EB neuropils; the latter is believed to maintain an internal compass of the fly. Through a series of queries and GUI operations, we construct this pathway as shown in Figure 3d. By combining connectivity and cell type information, it is possible to provide more information, such as tentative neurotransmitter type for each of the neurons involved in the pathway (see Video S7).

In light of the recent progress in understanding the brain structure of the larva [51], Neu-roNLP for the larva *Drosophila* is also provided (see Figure S2). Thus, it is possible now to compare larva and adult fly circuitry in, for example, the early olfactory system and mushroom body (see Video S8).

#### Building the Synaptome

Synapses can be added in NeuroNLP by using the information panel. For each pre- and post-synaptic partner, we provide a button to add into or remove from the workspace the synapses associated with the connection. In Figure 4a, we show the distribution of synapses from columnar neurons onto T4a neurons. This type of neurons have recently received major attention since they are the first neuron that exhibit a direction selective response to visual motion. By a few simple steps (see Video S9), it is possible to obtain a map of synapse onto T4a neurons similar to image provided by the data source [42], but with more powerful interactive 3D visualization. We can repeat the construction for other subtypes of T4 neurons (see Figure 4b-d).

**Figure 4:**
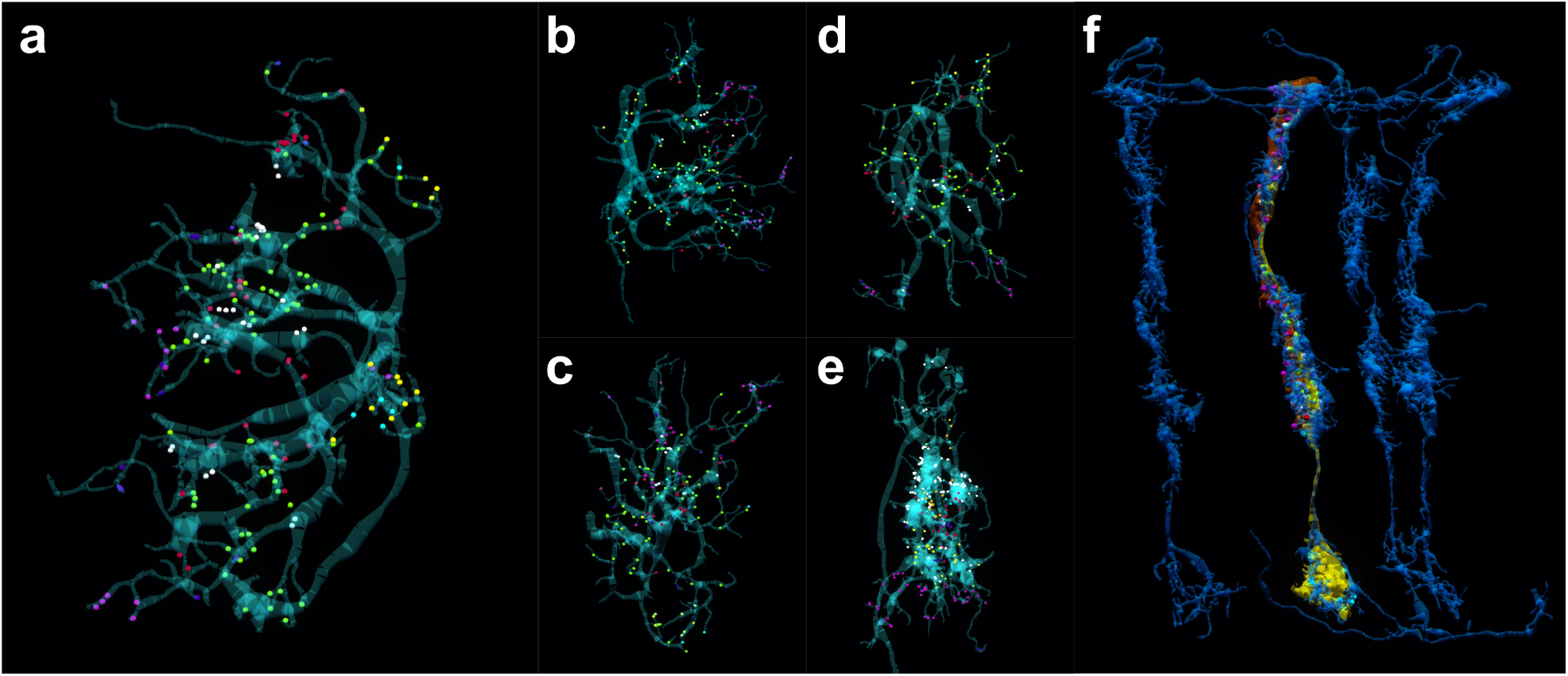
Using the synaptome map to probe the distribution of synapses. **(a-d)** Visualizing synaptic sites of inputs to four subtypes of T4 neuron, each are known to be sensitive to motion in one of the four cardinal directions. The 3D version can be accessed via tags: (a) “T4a_home_inputs” for the T4a neuron (see also Video S9), (b) “T4b_home_inputs” for the T4b neuron, (c) “T4c_home_inputs” for the T4c neuron and (d) T4d_home_inputs” for the T4d neuron. The T4 neurons are shown in transparent cyan. Synaptic sites are indicated by small spheres of different color. Each color correspond to synaptic input sites from a different cell type, but can be from multiple of such cells. (green) Mi1, (red) Tm3, (yellow) Mi4, (purple) Mi9, (cyan) C3, (dark blue) other T4 neurons, (white) presynaptic sites of this T4 neuron. **(e)** Distribution of pre- and post-synaptic sites of a Mi4 neuron in Medulla strata M2, M3 and M4. Majority of the inputs (shown in color dots) are located at the lower part of the dendrite in M2 and M3, while the terminals (white dots) are located at the upper part of the dendrite in M2. (green) inputs from Mi1, (purple) inputs from L5, (yellow) inputs from Mi9, (orange) inputs from R8, (blue) inputs from Dm2, (red) inputs from Dm4, (white) all outputs. **(f)** Distribution of synapses amongst a Dm9 neuron (blue) and an R7 (yellow) and an R8 (red) neurons. Sphere shows the location of synapses. (white) Dm9 to R8 synapses. (green) Dm9 to R7 synapses. (cyan) R7 to Dm9 synapses. (purple) R8 to Dm9 synapses. (red) R8 to R7 synapses. Access in NeuroNLP with tag color_input_medulla”.

In Figure 4e, we show the distribution of synaptic inputs and outputs of an Mi4 neuron in Medulla strata 2-5. In Figure 4f, the locations of synapse between pairs of R7, R8 and Dm9 neurons in the color vision pathway are visualized. However, to complete the synaptome information, it is necessary to also include data about neurotransmitter and receptor types for the current available synapse information. We expect that such data will be provided in the near future.

#### Building the Activity Map

In addition to providing a cell type map, a connectivity map, and the synaptome, NeuroNLP also provides an interface for accessing neurophysiology data to construct an activity map. Through NeuroNLP’s neurophysiology interface, users can search for available physiological data for a particular cell type, select datasets of interest and visualize them immediately in the web browser or download them in Neurodata Without Borders (NWB) format [52]. To our knowledge, this is the first instance of a completely searchable open physiology dataset for the fruit fly and we invite the research community to contribute additional fruit fly physiology data on the FFBO platform.

Figure 5 shows FFBO’s neurophysiology interface. Here, we query physiology data for an OR59b olfactory sensory neuron and for a photoreceptor, as shown in Figure 5a and Figure 5b, respectively. The queries result in lists of available data that users can scroll through and add to a list of dataset to be downloaded in the NWB format. Each column shows a particular type of experiment where the input waveform is shown on the top and the output at the bottom.

**Figure 5:**
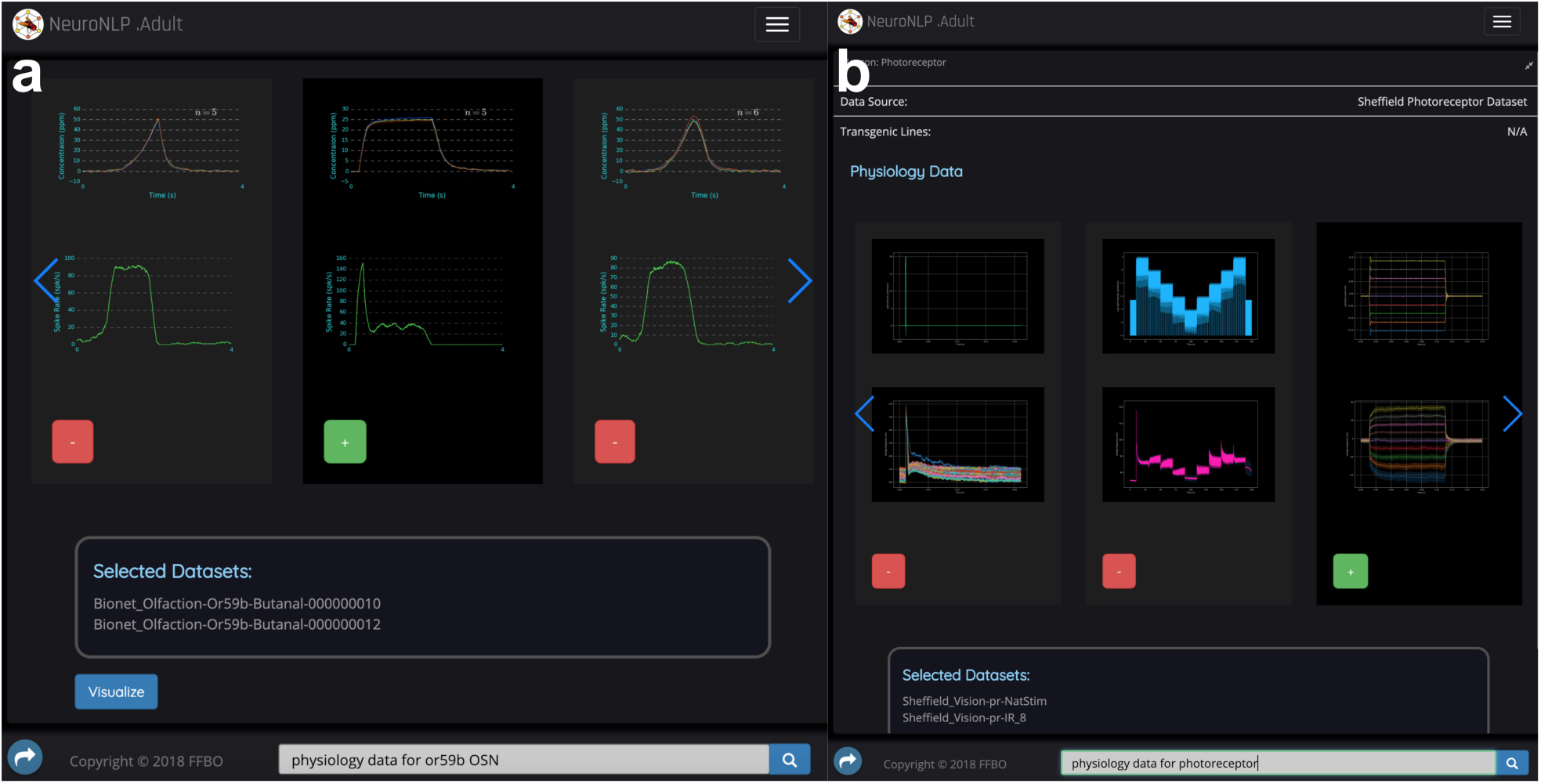
FFBO hosts an expanding activity map of the fruit fly brain, queryable in NeuroNLP. **(a)** Querying for physiology data for OS59b OSN recordings in a list of available data that users can scroll through and add to a list of dataset to be downloaded in NWB format. Each column shows a particular type of experiment where the input odorant waveform is shown on the top and the averaged spiking rate is shown at the bottom. The number of trials is indicated on the top. **(b)** Querying for physiology data for a photoreceptor.

### From Fruit Fly Brain Structure to Function with NeuroGFX

We present here a complete example of how NeuroGFX is used to construct the Central Complex [30]. Figure 6 shows the NeuroGFX interface for the CX. On the left panel, the biological structure of neuropils and neurons in the CX is visualized. On the right panel, NeuroGFX is loaded with an executable circuit diagram of the CX that reflects the biological data. The diagram is executable in that it is connected to a circuit model in the NeuroArch database that can be retrieved by Neurokernel for execution.

**Figure 6:**
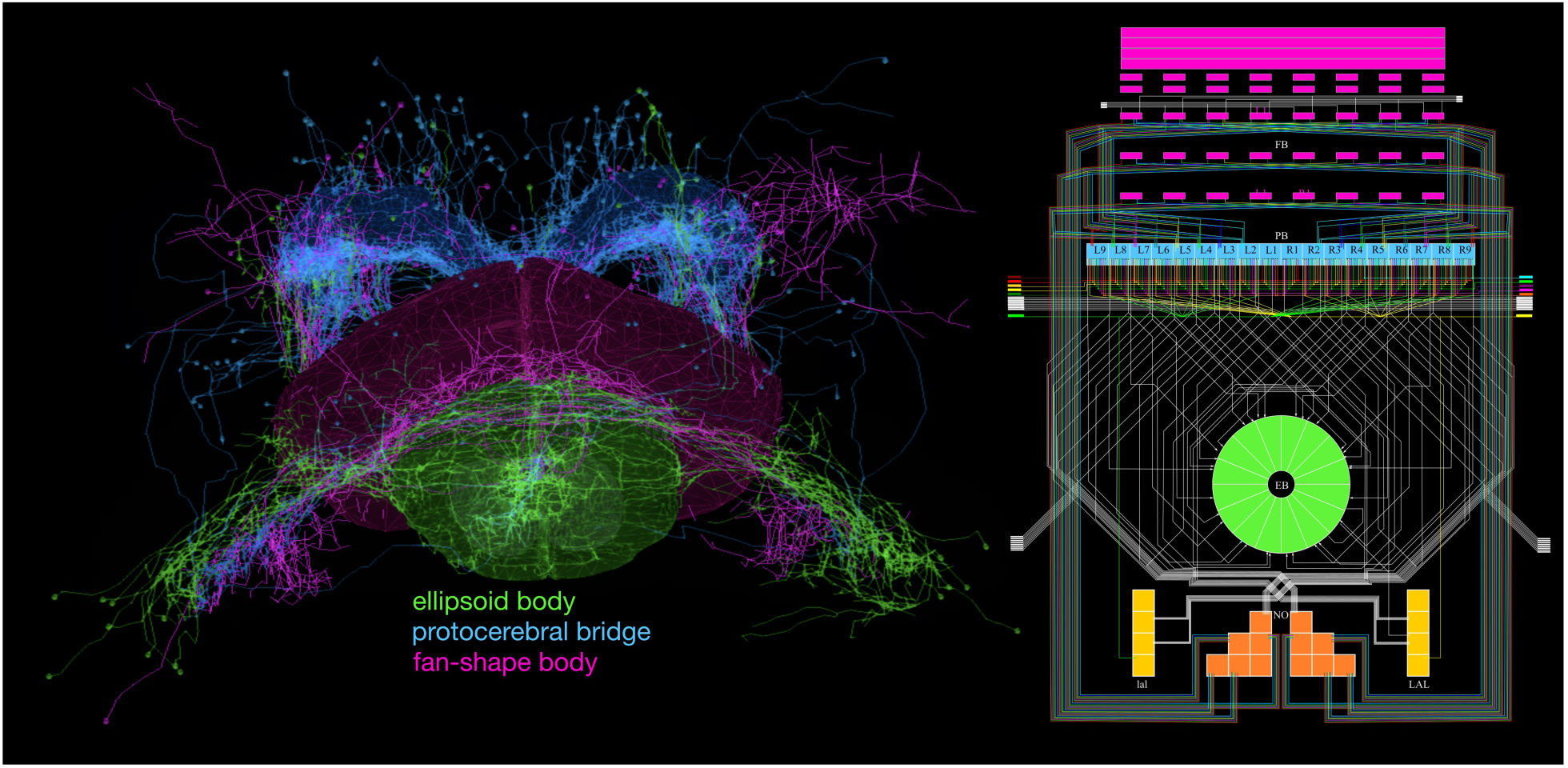
NeuroGFX enables the exploration of circuit function from circuit structure. A neuropil level exploration of the Central Complex (CX). The circuit structure of the CX, including neuropils and neurons in the CX (left), are visualized in parallel with the executable circuit diagram of CX (right). NeuroGFX loads the circuit model associated with the diagram from the NeuroArch database. Its GUI enables users to remove/add neurons in the circuit by clicking on the neurons in the left panel or the circuit diagram in the right panel. Reconfiguration of the circuit is then done by the NeuroArch database, and the resulting executable circuit is retrieved by Neurokernel for execution.

On the local circuit level, NeuroGFX supports the exploration of the function of circuits of manageable size, *e.g.*, a canonical circuit or a basic building block in an LPU. In Video S10, we construct a cartridge of the Lamina neuropil. The complete shape of neurons and connectivity between neurons in a single cartridge of the lamina have been determined by serial electron microscopy [13]. Using NeuroGFX, we visualize side by side both the morphology (skeletons) of the neuron and the circuit diagram.

The neural circuit configuration, manipulation and composition capabilities in NeuroGFX are a significant step towards providing *design automation* tools for computational neuroscience. We have already used these tools to develop and demonstrate computational models for various subregions of the fruit fly brain including retina, lamina, antenna and antennal lobe. Different implementations of the retina, independently developed [29, 53], can be interfaced with the lamina, validating the composibility of LPU implementations enabled by the design of the Neurokernel architecture [41].

## Discussion

Recent tools in fly neurogenetics and neurobiology have unveiled a staggering and constantly increasing amount of structural fruit fly brain data. While lagging behind, and thanks to increasing massively parallel computing power, computational studies have focussed on the emulation of the function of brain circuits. The link between the two has been largely lacking, however.

The FFBO raises to this challenge by a tight integration of the neurogenetics, anatomy and physiology data with computational modeling data of the fruit fly brain into a single database. By endowing the FFBO with a powerful query, circuit execution and visualization framework, we have taken a key step towards creating a platform for exploring and validating brain function from structure. Among others, FFBO enables the comparison of different computational models of the same brain circuit or reused circuit motifs as well as the comparison between the function of analogous circuits in the adult and larva fly.

FFBO greatly increases the accessibility of fruit fly brain data, and provides tools for creating structural circuits of interest. It also facilitates disseminating results of complex queries through the use of tags, uniquely enabling a collaborative exploration and dissemination of neural circuit compositions. Furthermore, FFBO provides neurophysiology data in public domain, a first step towards building an activity map of the fly brain and for studying the function of neural circuits and their biological validation. Finally, FFBO enables the study of brain function on multiple levels of abstractions including the circuit and whole brain level.

To meet the increase of future data [17], the underlying design of the main FFBO components is scalable. For example, the NeuroNLP interface and its underlying NeuroArch database are easily extensible with new data types and can handle large amounts of additional data. Support for an *in silico* experimental workbench and for *in silico* optogenetic experiments can be added as well.

For neurobiologists and neurogenetists, the tools we created will catalyze the discovery of fly circuit function by creating a reservoir of neurons and neurons types that can be easily targeted using genetics. For computational/theoretical neuroscientists, the tools will allow them to transcend the physical limitations of biological experiments that generate massive amounts of data. For computer scientists/engineers new architectures for deep learning networks can be explored. Model libraries built by computational/theoretical neuroscientists and computer science/engineers will, in turn, enable easy to configure computational experiments for neurogeneticists/neurobiologists to test various hypotheses. With these tools, thereby, the FFBO ecosystem will accelerate the pace of creating computational models of fly brain circuits, and of uncovering the logic of neuroinformation processing in the fly brain.

Finally, we note that in order to create accurate large scale models of the brain of an organism, independent labs must be able to easily share and integrate independently created computational models. The standardized model of communication among LPUs makes such an integration under FFBO a reality. Furthermore, by adopting the standards defined by Neurodata Without Borders (NWB) [52] FFBO enables users to upload data to NeuroArch in the NWB format. Note that a circuit model format has not yet been adopted in the literature.

The key features of the FFBO differ in fundamental ways from other platforms such as the Virtual Fly Brain [23] and the Insect Brain Database (https://insectbraindb.org). While the Virtual Fly Brain and Insect Brain Database mainly serve confocal imaging stacks, they do not provide any electrophysiology and imaging data. The point and click interface they provide limits the comprehensiveness of the queries and the resulting data to be used in designing and performing new experiments. This is to be contrasted with the simpler to use and expression rich natural language interface provided by FFBO. More importantly, both repositories also lack native support for integration of executable circuit models and, anatomical and physiological data, i.e., these platforms lack tools for bridging structural data and functional data.

By supporting the exploration of new neural pathways, new circuit diagrams and new executable circuit models, FFBO has become a key resource for exploring the function of neural circuits from structured data. The availability and easy of use of these capabilities have raised a number of important new questions regarding the functional map of fruit fly brain circuits. First, with the natural language processing interface of NeuroNLP we demonstrated the visual display of known neural pathways. Can FFBO support the exploration of novel pathways between an arbitrary pair of neuropils, say the optic lobes and the central complex (see Figure 3)? Second, can the data shown in Figure 5 be displayed in conjunction with a display of the putative neurons the recordings may be associated with? Third, can FFBO support NeuroCAD utilities (see Figure 1) that enable the design of neural circuits akin to a Computer Aided Design tool for silicon circuits? Fourth, the executable circuit in Figure 6 was designed and implemented by hand. An arbitrarily chosen subcircuit in the same figure can be executed and functionally evaluated. Can arbitrary brain circuits be constructed, executed and biologically validated?

The answer to all these questions is explored with the interactive open computing platform FlyBrainLab, currently under development [54]. Users are encouraged to download and contribute to its development. The results will be published elsewhere.

There are other efforts in the literature that aim at providing data repositories and software tools for model organisms. Two major efforts stand out: the OpenWorm (http://openworm.org) and the Allen Brain Atlas (http://brain-map.org). The OpenWorm project is dedicated to creating a virtual *C. elegans* in computer program with all features of its behaviors, by crowdsourcing a community of computational neuroscientists and computer scientists across the world [55]. The Allen Brain Atlas is a repertoire of brain data with an emphasis on mouse as a model organism. Unlike the OpenWorm project driven by the open source community, the Allen Brain Atlas is an in-house effort by an industrial-scale team of researchers from the Allen Institute. It serves different stakeholders with various data repositories, including the Allen Brain Observatory hosting in-vivo recording, the Cell Type Database containing a survey of biological features, the Allen Software Development Kit for data analysis and model simulation, and many other datasets and toolkits [56]. In alignment with the OpenWorm project and the Allen Brain Atlas, FFBO champions the focus on model organism in quest of understanding brain functions.

## Methods

### System Architecture of the Fruit Fly Brain Observatory

The system architecture of the FFBO consists of three levels of abstraction, as shown in Figure S1. The lowest level consists of the (i) FFBO Processor operating as central control for registering services and routing messages (ii) NeuroArch Database and associated APIs for querying and manipulating the data, and (iii) Neurokernel Engine for the emulation of *Drosophila* brain circuits on massively parallel GPU clusters. The mid level consists of the NLP Module, a natural language translator for Neuroarch API access, a Visualization Engine for anatomy and physiology data and, a Neural Circuit Design Module, a set of tools for exploring brain circuit models. Finally, at the top layer, reside the NeuroNLP and NeuroGFX front-ends.

The messaging between components is handled using the Web Application Messaging Protocol (WAMP) (https://wamp-proto.org), stable implementations of which exist for all major languages. The FFBO Processor serves as a WAMP router, and components providing services register their services with the FFBO Processor. Applications then call these services using Remote Procedure Calls (RPCs). Multiple backend components can register with the processor. For example, when multiple NeuroArch components launched on different machines are registered, different users can be served on a local machine for better user experience and redundancy.

Finally, all backend components are containerized using Docker (https://www.docker.com). This allows for easy sharing, installation and deployment, and for compartmentalization of the distributed components.

### NeuroArch Database

The NeuroArch data model preserves the structural and semantic relationships between different biological and modeling objects. Its query interface provides an object-graph mapping (OGM) that enables both neurobiologists and neural circuit model designers to easily perform sophisticated queries relevant to their respective needs without having to explicitly specify complex query strings. OGM provides methods associated with each object that dynamically construct and execute queries. Two key differences between NeuroArch’s OGM and that of currently available general purpose OGMs are (i) its use of the hierarchical data model to enable extraction of subcircuits owned by nodes corresponding to specific subdivisions of biological components or circuit abstractions, and (ii) its ability to use the subgraph extracted by an OGM query as the starting point for traversals by subsequent queries or as an operand that may be passed to graph operators. Thus queries can be composed with other queries using various operators. For example, query results can be added (union) or subtracted (difference), and thereby, novel, more complex, queries easily constructed.

The Neuroarch API is exposed through a Remote Procedure Call (RPC) initiated by the NeuroArch component; the latter component accepts a JavaScript Object Notation (JSON) object. The JSON object specifies the method/operator, arguments for the method/operator and the object or class to access the method/operator. Multiple access methods may be specified in a single RPC call; an RPC specific results queue allows intermediate results as the starting point or as operands for subsequent method/operator calls. Additionally, the NeuroArch component maintains a user specific memory queue, allowing for subsequent queries to act on present or past results.

Technical documentation and code for NeuroArch and NeuroArch component is available on Github. Further information on the design considerations for NeuroArch data model and API are described in Neurokernel RFC #5 [40].

### Neurokernel Engine

Neurokernel is an open-source engine implemented in Python for the collaborative emulation and validation of fruit fly brain models on multiple Graphics Processing Units (GPUs). Neurokernel defines communication interfaces that specify how spikes and neuron membrane states are transmitted between LPU models. It provides the necessary machinery for efficient transfer of information between different LPU implementations. Other than the communication API, it does not impose any other restrictions on the LPU implementation.

Technical documentation and the code is available on Github. Further information about the API design and the implementation performance can be found in [41].

### Natural Language Processing Module

To translate natural language queries from NeuroNLP into machine code understandable by the NeuroArch API, the current NLP module is built upon the Quepy framework (https://github.com/machinalis/quepy). The NLP Module attempts to generate a semantic parse tree from the natural language query, optimizes it where possible, and finally produces the NeuroArch API representation from the optimized parse tree. If it fails to generate a semantic parse tree, it informs the requestor that it could not interpret the query. Documentation and code is available on the Github repository (https://github.com/fruitflybrain/neuroarch_nlp).

### NeuroNLP Demos

An easy-to-use scripting tool supports the creation of guided demonstrations that can be composed by defining an ordered list of queries, user interactions and informational messages to be displayed during the demonstration. We employ this functionality to provide demos that help users to get started with NeuroNLP, and it can easily be used to provide demos for specific educational or research dissemination purposes. The demo scripts are formatted in JavaScript Object Notation (JSON). An example is provided as following:

~~~
“demo1”: {
    “script”: [
    [“notify”, {“message”: “Querying for Mi1 neurons”,
                              “pause”: 1000, “timeout”: 2000}],
    [“search”, {“query”: “Show Mi1 neurons”}],
    [“click”, {“label”: “Mi1-home”]
  ]
}
~~~

### NeuroNLP Tags

The results obtained at any point of query, possibly through multiple user interactions, can be shared easily using the ‘tag’ feature, which can exactly recreate the state of the sharer’s session on any modern browser, including smartphones, when shared as a link to the public and/or collaborators. A tag consists of a complete snapshot of the current workspace; loading the tag on any other device will recreate the same workspace, including camera angles and neuron colors. Therefore, The tag feature can be used for easy sharing of circuits, for recreation of figures from published papers or for use directly as 3D interactive figures in publication, presentation and education.

### Information Panel in NeuroNLP

By clicking on a neuron displayed in the main workspace of NeuroNLP, data associated with the neuron are shown in the information panel. For neurons originated by the FlyCircuit database, we display a list including the GAL4 driver used, putative neurotransmitter, gender and age, putative birth time and soma coordinate. In addition, we show the confocal images of the neuron. These data are directly provided by the FlyCircuit database. When available, we also include a link to the neuron in Virtual Fly Brain. For neurons obtained using EM methods, we only list the neurotransmitter information if they are available in the literature. The information panel also contains the synaptic profile of the neuron. For neurons from FlyCircuit database, a bar chart is provided to summarize the percentage of putative pre- and post-synaptic partners of each driver line. A list of pre-synaptic and post-synaptic neurons is at users’ full disposal, showing the number of putative synaptic contacts. Users can click on the ‘+’ button to add a neuron individually to the workspace. For EM-based data, since precise number of synapses is available, they are listed in the pre-synaptic and post-synaptic neurons. The bar chart is again used to summarize percentage of each type of neuron that are pre- and post-synaptic to the neuron in question. Moreover, for dataset in which information about individual synapses is available, the information panel provides a button to visualize all the synapses in their respective position between the pre- and post-synaptic neurons.

### Inferred Synaptic Connectivity

Connectivity of neurons in the FlyCircuit database are inferred via the method detailed in [25]. The dendritic and axonal terminals of each neuron are inferred by the SPIN method detailed in [57].

### Visualization Engine

The visualization engine processes the anatomical data retrieved from NeuroArch and renders the data in 3D in the web browsers at the user end. Two data formats are supported by the engine: (i) the mesh format that represents a neuropil as a 3D surface consisting of vertices and triangular faces, and (ii) the SWC format that stores a neuron as a 3D tree with vertices and edges. Each tree vertex is labeled with soma, dendrites or other anatomical properties.

The visualization engine is built upon three.js (https://threejs.org), a cross-browser JavaScript library for creating and displaying 3D computer graphics in web browsers. The visualization engine provides a user with basic operations such as translation and rotation as well as advanced features including changing colors and highlighting neurons.

To allow researchers to share customized neural circuits of interest among other researchers, the API of the visualization engine supports importing and exporting the configuration of a customized visualization from and to NeuroArch. This allows users to visualize the same neural circuits with exactly the same configuration.

### NeuroGFX User Interface

Circuit diagrams in the NeuroGFX are created by hand in Scalable Vector Graphics (SVG) format. They are loaded in NeuroGFX at a corresponding level of abstraction and bound to biological structure by a custom javascript program. The whole brain level diagram of the network of Local Processing Units (LPUs), i.e., model abstractions of neuropils, is laid out based on [58]. Through the interface, the circuit diagram can be reconfigured to include any subset of neuropils, allowing to probe how a particular composition of neuropils leads to a specific brain function. On the neuropil level, NeuroGFX allows users to study the I/O of each LPU. The circuit diagram of the CX is created according to [30]. On the local circuit level, the focus of NeuroGFX is to explore the function of a circuit of manageable size, e.g., a canonical circuit in an LPU, thereby facilitating the understanding of the basic building blocks of their respective LPUs. The complete shape of neurons and connectivity between neurons in a single cartridge of the lamina have been determined by serial electron microscopy [13]. The circuit diagram of the cartridge is created according to the connectome data. By clicking on the Load Cartridge button, NeuroGFX is instructed to fire a series of queries to the NeuroArch Database where a model of the retina and lamina network resides. Upon retrieving the circuit model, information about an individual neuron can be shown by hovering the mouse over the neuron in the circuit diagram. When the Open NK button is clicked, the Neurokernel Engine is instructed to retrieve the configured circuit from the NeuroArch Database for execution.

### Neural Circuit Design Module

To facilitate interaction with circuit diagrams in NeuroGFX, the Neural Circuit Design Module offers a set of tools for manipulating neural circuits in the scalable vector graphics (SVG) format. To enable seamless integration between the circuit diagram and the 3D visualization of a neural circuit, additional attributes in the SVG file associate an object in the diagram with the name of a neuron or with the property of a synapse. The object based on JavaScript code can then be manipulated in the browser.

### Code Availability and Current Hosting

The source-code for the Fruit Fly Brain Observatory is publicly available on Github under the account https://github.com/fruitflybrain. The FFBO is also publicly accessible at http://fruitflybrain.org using any modern web browsers, including those running on smartphones. Core components of the FFBO have been released in a containerized form to be easily installed and launched locally. In addition, a repository available at https://github.com/fruitflybrain/ffbo.launcher can be used to connect individual components such that any lab can easily host all FFBO services locally, to access their own data, and independently develop interoperable network models. We also provide an Amazon Machine Image to allow users to launch a full copy of the FFBO services on Amazon Web Services EC2 with a few simple clicks.

## Supporting information

Sharing a neural circuit using tags in NeuroNLP.

Demo player constructs a neural circuit step by step from a demo script.

Building a cell type map using NeuroNLP (1).

Building a cell type map using NeuroNLP (2).

Building a cell type map using NeuroNLP (3).

Constructing the visual motion pathway from connectome data.

Constructing a visual pathway to EB.

Comparison of early olfactory system and mushroom body between adult and larva Drosophila using NeuroNLP.adult and NeuroNLP.larva.

Visualizing the synapses of T4 neurons.

Local circuit exploration on NeuroGFX.

## Acknowledgements

The “Fruit Fly Observatory” was selected as a Phase I winner of the 2016 Open Science Prize Challenge and was initially supported in part by the NIH, Wellcome Trust and HHMI. The research reported here was also supported, in part, by NSF under grant #1544383, in part by BBSRC #BB/M025527/1, in part by AFOSR under grant #FA9550-16-1-0410 and in part by the Higher Education Sprout Project funded by the Ministry of Science and Technology and Ministry of Education in Taiwan. The authors would like to thank Jonathan N. Martin for contributing to FFBO installation and launcher scripts.

## Author Contributions

A.A.L., A.-S.C., D.C., C.-C.L., P.R. conceived of the study. N.H.U., C.-H.Y., A.T. and Y.Z. created prototype of the open-source platform, and improved it together with M.K.T. and T.K.L.. Y.-C.H., C.T.W., N.H.U, C.-H.Y. Y.Z. C.L.O. processed data. N.H.U., A.T., C.-H.Y. Y.Z., C.L.O., D.F. contributed to computational modeling. N.H.U., C.H.Y., A.T. Y.Z., D.C., and A.A.L. wrote the manuscript with input from the other authors.

## Supplementary Material

**Supplementary Figure S1:**
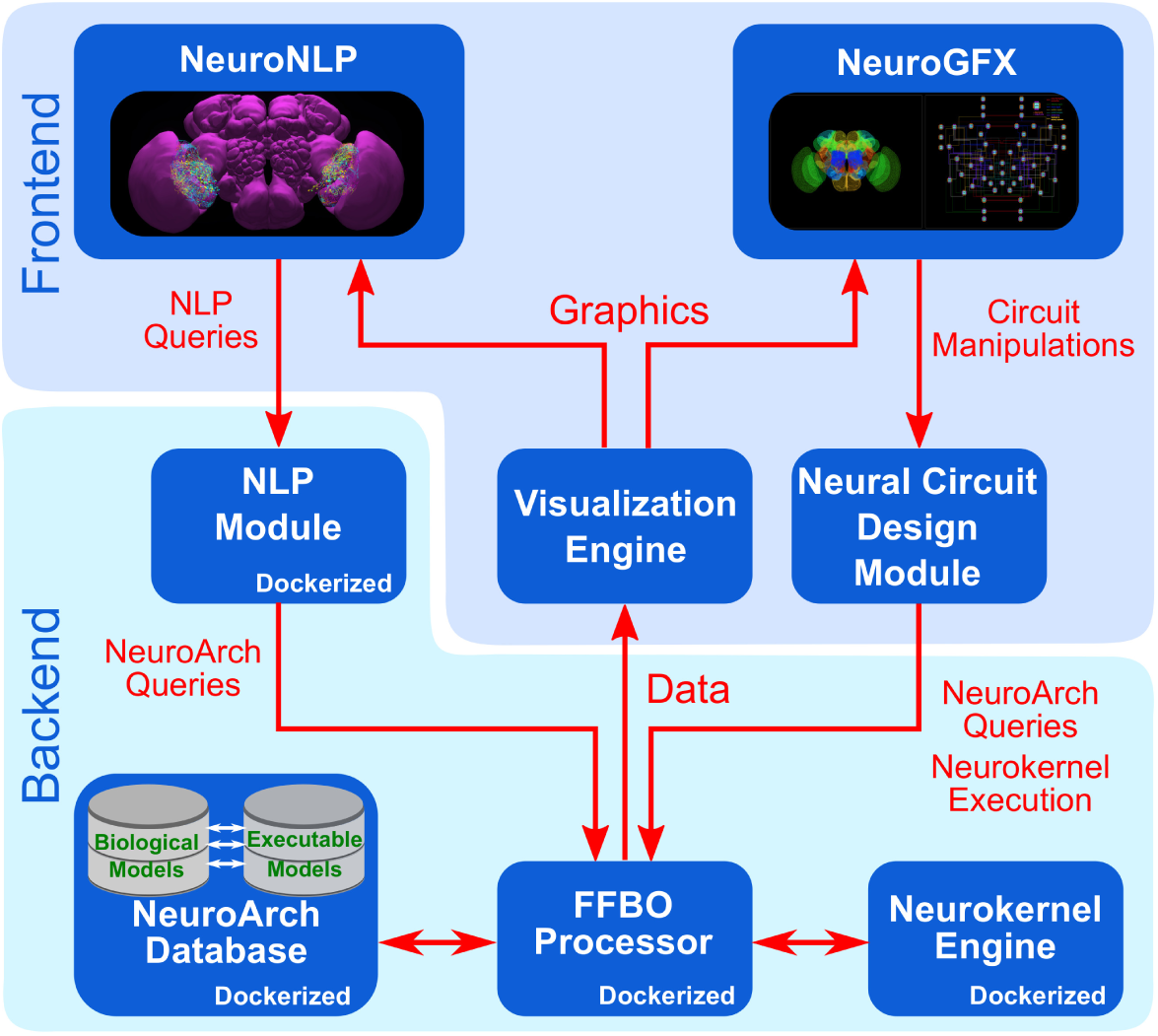
The modular system architecture of the FFBO. The FFBO provides 2 user level applications, the NeuroNLP and NeuroGFX. Supporting these frontend applications are backend services provided by an FFBO processor connected to the NeuroArch data service server, and the Neurokernel execution service server. An NLP module is included to handle the parsing of plain English queries. A visualization engine is used for visualization of biological data in the frontend, respectively. The backend components are all containerized using Docker, making replacement of each component and using of multiple components straightforward.

The message bar (in light blue in the videos) provide narratives of the videos and will help the reader understand the steps being taken and potential scientific value of the results.

**Supplementary Figure S2:**
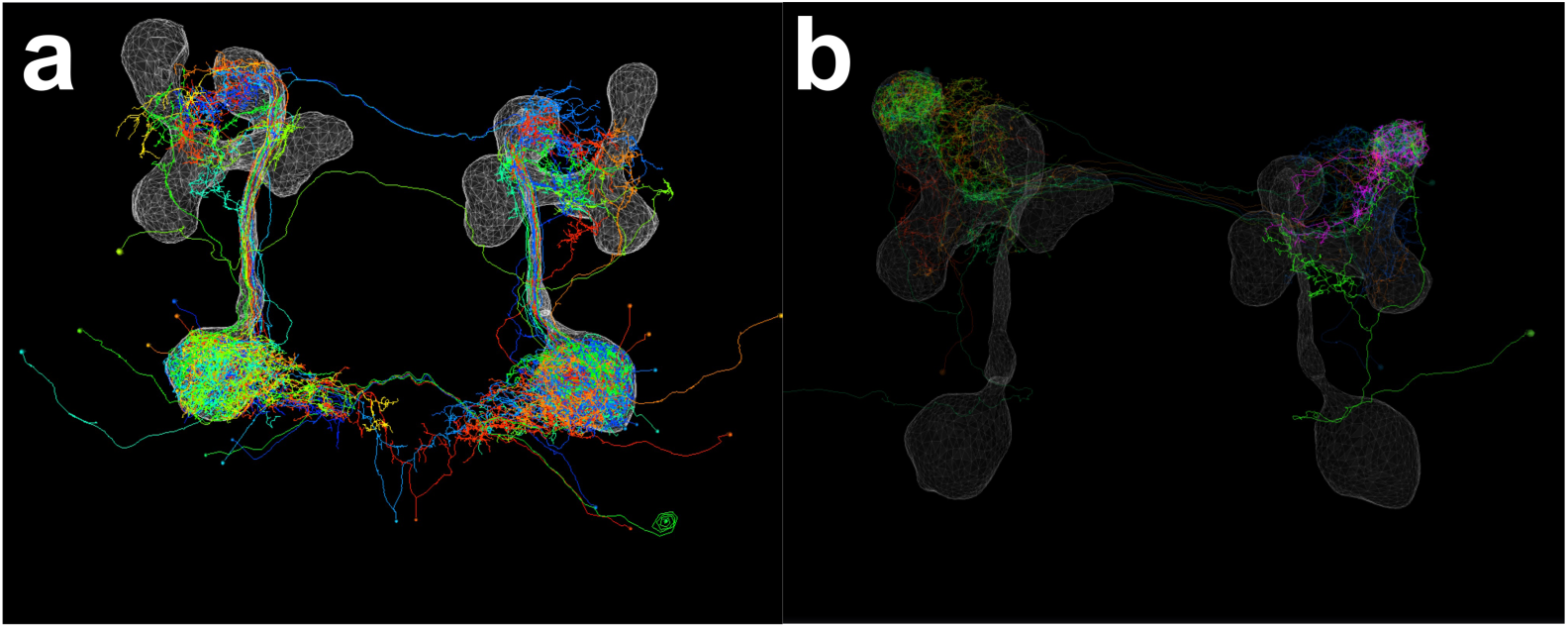
NeuroNLP Larva currently provides data from [43] and [44]. Each of the figures in these papers is associated with tags. **(a)** Retrieved tag for Figure 4a in [43]. **(b)** Retrieved tag for Extended Figure 5 UVL in [44]. These tags provide interactive 3D versions of the static figures.

**Supplementary Video S1:** Sharing a neural circuit using tags in NeuroNLP. This video provides an example of retrieving a tag to obtain the exact 3D visualization of an neural circuit as stored by the creator of the tag.

**Supplementary Video S2:** Demo player constructs a neural circuit step by step from a demo script. This video shows how a demo of circuit query can be automated using the demo player.

**Supplementary Video S3:** (related to Figure 2a) Building a cell type map using NeuroNLP. Direct querying of Mi1, Mi4, Mi9, Dm8 and Dm2 neurons in specific columns of the Medulla.

**Supplementary Video S4:** (related to Figure 2b) Building a cell type map using NeuroNLP. Determining subclasses of LPTCs in the fruit fly. More than 20 anatomical classes of LPTCs are known in the blowfly (Hausen, 1984). Little is known about the existence and if it exists, the function of their homologs in the fruit fly visual system. Here we use NeuroNLP to identify LPTCs that may exist in the FlyCircuit data, and their anatomical classes.

**Supplementary Video S5:** (related to Figure 2c) Building a cell type map using NeuroNLP. Discovering new Lobula Plate local neurons to drive new experiments and computational studies. Glutamatergic inhibitory local neurons that bi-stratify in layers 3/4 of Lobula Plate (LPi neurons) has been identified [49]. We query in NeuroNLP to discover a few more types of LPi neurons.

**Supplementary Video S6:** (related to Figure 3a-c) Constructing the visual motion pathway from connectome data. The motion detection pathway is one of the most studied circuits in *Drosophila* [28]. Motion information is processed in two parallel pathways, with motion sensitive neurons in the L1 pathway processing motion associated with brightness increments (ON pathway) while the L2 pathway processes brightness decrements. T4 neurons are the first neurons in the L1 pathway whose responses are direction selective [59, 60]. Columnar neurons involved in processing the motion information in a retinotopic fashion have been studied extensively, but the role of the non-columnar neurons in processing motion information remains obscure. In this video, we use NeuroNLP to explore direct and indirect inputs to a T4 neuron.

**Supplementary Video S7:** (related to Figure 3d) Constructing a visual pathway to EB. In [50], a two-stage visual pathway between the medulla and the bulb (BU, or lateral triangle) was uncovered. This identifies potential sources of visual input to ring neurons that links BU and EB, a neuropil that is believed to maintain an internal compass of the fly. We can inspect this pathway by a few queries and GUI operations.

**Supplementary Video S8:** Comparison of early olfactory system and mushroom body between adult and larva *Drosophila* using NeuroNLP.adult and NeuroNLP.larva.

**Supplementary Video S9:** (related to Figure 4a) Visualizing the synapses of T4 neurons. When available, synaptic sites can be easily visualized in NeuroNLP. We focus here on the distribution of synaptic inputs onto a T4a neuron from its major inputs [42]. The video provides all the steps for creating a synaptome for T4 in NeuroNLP.

**Supplementary Video S10:** Local circuit exploration on NeuroGFX. Execution of a cartridge in the Lamina. We constructed an example NeuroGFX for the Lamina cartridges. This can be accessed in https://neurogfx.fruitflybrain.org by double clicking on the “LAM” LPU, and then double clicking on any of the cartridges when the lamina page is brought up. The circuit diagram is drawn based on EM data [13]. By clicking on the “Load Cartridge” button, the cartridge model will be constructed from queries retrieving data from the NeuroArch database. This allows users to examine model parameters for each of the neurons by hovering the mouse over the corresponding block. A user can then click on any of the neurons to disable/ablate or re-enable it. The disabled neuron will be removed from the neuron skeleton window, as well. After reconfiguring the neural circuit by these simple clicks, the user can execute it by clicking on the “Open NK” button. This fires up additional NeuroArch queries to remove the disabled neurons from the previous query result of the full cartridge. Neurokernel will then retrieve the latest, modified cartridge circuit from NeuroArch and execute the circuit. After circuit execution finishes, the execution results are transferred to the browser for visualization. Here the top row shows the input waveform of the photoreceptors, and the response of all the neurons are shown in the plot at the bottom. An animated visualization can be played by clicking on the play button at the bottom left corner.

